# Alternate Grainy head isoforms regulate *Drosophila* midgut intestinal stem cell differentiation

**DOI:** 10.1101/2020.12.20.423699

**Authors:** Nicole Dominado, Franca Casagranda, James Heaney, Nicole A. Siddall, Helen E. Abud, Gary R. Hime

## Abstract

Regeneration of the *Drosophila* midgut epithelium depends upon differential expression of transcription factors in intestinal stem cells and their progeny. The *grainy head* locus produces multiple splice forms that result in production of two classes of transcription factor, designated Grh.O and Grh.N. *grainy head* expression is associated with epithelial tissue and has roles in epidermal development and regeneration but had not been examined for a function in the midgut epithelium. Null mutant clones had a limited effect on intestinal stem cell (ISC) maintenance and proliferation, but specific loss of the Grh.O isoform results in loss of ISCs from the epithelium. This was confirmed by generation of a new Grh.O mutant to control for genetic background effects. Grh.O mutant ISCs were not lost due to cell death but were forced to differentiate. Ectopic expression of the Grh.N isoform also resulted in ISC differentiation suggesting that the two isoforms act in an opposing manner. Grh.O expression must be tightly regulated as high-level ectopic expression in enteroblasts, but not ISCs, resulted in cells with confused identity and promoted excess proliferation in the epithelium. Thus, midgut regeneration is not only dependent upon signalling pathways that regulate transcription factor expression, but also upon regulated mRNA splicing of these genes.

## Introduction

Epithelial tissues provide barrier function between organs and their surrounds and must be able to repair or replace damaged cells (1). The *Drosophila* midgut epithelium contains two types of differentiated cells, the absorptive enterocyte (EC) and enteroendocrine (EE) cells and is an excellent model to study the molecular processes that regulate maintenance of epithelial stem cells and their differentiation. ECs are large polyploid cells that make up the bulk of the midgut epithelia, interspersed with hormone producing EE cells, which regulate peristalsis (2), cell fate (3, 4) and intestinal stem cell (ISC) proliferation (5, 6). The differentiated cells are constantly replaced by a population of regenerating ISCs scattered along the basal surface of the epithelium. ISC divisions produce daughter ISC and transient enteroblast (EB) progenitors. EBs can differentiate into either an absorptive EC or a secretory EE (7–9) although data indicate that ECs and EEs arise from different progenitors and EE cells can directly originate from ISCs primed to form EE cells (3, 4, 10, 11). The Grainy head (Grh) family of transcriptional regulators is conserved across metazoan lineages and factors in epidermal barrier formation, wound healing, tubulogenesis and cancer. The initial presence of the *grainy head* gene family coincides with the evolution of epithelia, highlighting the importance of this family for epithelial regulation (12).

*Drosophila* has a solitary *grh* gene whereas mammals have evolved three *Grhl* genes, *Grhl-1, Grhl-2,* and *Grhl-3.* Studies in both *Drosophila* and mice have demonstrated that specific Grh proteins are essential for formation and maintenance of epithelial tissues. Vertebrate studies have produced complex results with different members of the Grh family associated with induction of differentiation and in some cancer studies, stemness (13–17).

In *Drosophila,* the single *grh* (also known as *Elf-1* / *NTF-1)* gene is alternatively spliced. *grh* transcripts can be classed into two groups with those containing exons 4 and 5 known as O-isoforms (Grh.O and Grh.O’) whereas transcripts without these exons are classed as N-isoforms (Grh.N and Grh.N’) (18) (Figure 1A). O-isoforms have been reported as being restricted to neural tissues in third instar larvae with N-isoforms associated with epithelial maintenance and wound repair in non-neural tissues (18–21). Analysis of isoform specific functions of Grh in the midgut would facilitate an understanding of how this gene family may play important roles in regenerative epithelia. Here we show that both isoforms are expressed at very low levels in the midgut epithelium where they play different roles in regulating ISC maintenance, proliferation and differentiation.

**Figure 1.**
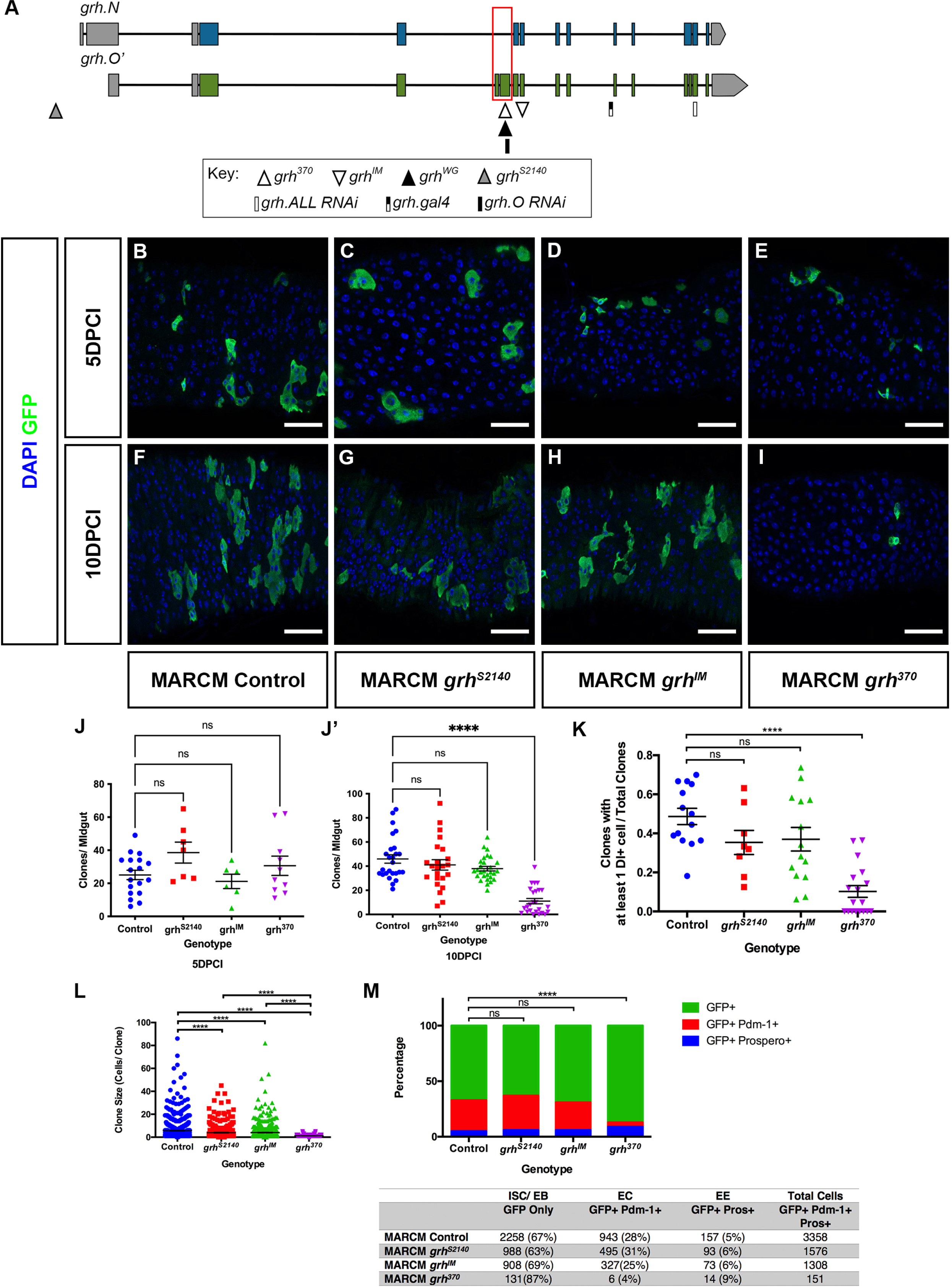
*grh* O-isoforms maintain ISC identity. **A)** Schematic of *grh* mRNA transcripts. The single *grh* gene is alternatively spliced to produce two classes of transcripts based on the splicing of exons 4 and 5 (red box). Transcripts with exons 4 and 5 are classed as O-isoforms while those without these exons are classed as N-isoforms. *grh.N* and *grh.O’* are shown as representatives of the two classes. Positions of mutations are indicated by triangles and the grh.Gal4 insertion by a black and white bar. Sequences that were utilised for RNAi constructs are indicated by bars. **B-I)** Representative maximal intensity z-projections of GFP (green) marked control, *grh*^*S2140*^*, grh*^*IM*^ *and grh*^*370*^ MARCM clones at 5PDCI and 10DPCI. Scale Bar 40μm. **J-J’)** Quantification of the mean number of clones per midgut at 5 and 10 days post clone induction (DPCI). MARCM control (5DPCI: n=18 midguts, 10DPCI n=26 midguts) and *grh* null (*grh*^*S2140*^ – 5DPCI: n=7 midguts, 10DPCI n= 23 midguts; *grh*^*IM*^ – 5DPCI: n=6 midguts 10DPCI: n=28 midguts) O-specific mutant *grh*^*370*^ (5DPCI: n= 10 midguts, 10DPCI: n= 25 midguts) clones are lost at 10DPCI. (Mean ± SEM, One-way ANOVA with Dunnett’s test, ns= not significant, ****p<0.0001). **K)** Proportion of MARCM clones at 10DPCI containing at least a single Dl+ ISC in control (n=14), *grh*^*S2140*^ (n=8), *grh*^*IM*^ (n=14) and *grh*^*370*^ (n=18) midguts. In comparison to control clones, there is a significant decrease in the fraction of *grh*^*370*^ clones containing ISCs. (Mean ± SEM, One-Way ANOVA with Tukey’s test, ns= not significant, ****p<0.0001). **L)** *grh*^*370*^ MARCM clones affect ISC proliferation. Control clones are larger in size compared to *grh* mutant clones at 10DPCI. Clonal size was calculated by counting the number of cells within each clone in control and *grh* mutants. Control – n= 770 clones, 26 midguts. *grh*^*S2140*^ – n= 526 clones, 15 midguts. *grh*^*IM*^- n=554 clones, 21 midguts. *grh*^*370*^ – n=269 clones, 25 midguts. (Mean ± SEM, One-Way ANOVA with Tukey’s Test, ***p=0.0002, ****p<0.0001). **M)** *grh*^*370*^ MARCM clones affect ISC differentiation. Quantification of the cellular composition of MARCM clones immunostained with Prospero and Pdm-1. The number of ECs (GFP+ Pdm-1+), EE cells (GFP+ Prospero+) and ISC/EBs (GFP+) for control and *grh* mutant clones was quantified for each genotype and percentage values calculated. (χ^2^ Test, control vs. *grh*^*S2140*^ p=0.8325, control vs *grh*^*IM*^ p=0.8650 and control vs. *grh*^*370*^ ****p<0.0001).

## Results

### Grainyhead O-isoforms maintain ISCs by preventing differentiation

We examined two *grh* amorphic (null) alleles, *grh*^*S2140*^ and *grh*^*IM*^, in addition to a hypomorphic allele, *grh*^*370*^ which only disrupts function of all O-isoforms (18). Homozygotes of each allele are embryonic lethal, hence the MARCM system (22) was utilized to generate GFP marked homozygous *grh* mutant clones in posterior midgut tissue.

Initial analysis of clones at 5 and 10 days post clonal induction (DPCI) revealed that by 10 DPCI the number of *grh*^*370*^ clones per midgut was less than that of control and *grh* null clones (Figure 1B-J) at 10DPCI. Given that clones arise and are maintained by ISCs, a loss of GFP marked clones at the later time point would suggest that Grh O-isoforms are required for the maintenance of ISCs. To further investigate this hypothesis, control and *grh* clones 10DPCI were immunostained with the ISC marker, Delta (Dl). In comparison to controls (0.49±0.04 clones), there was a slight but non-significant decrease in the proportion of clones with at least 1 Dl+ cell in *grh* null mutants (*grh^S2140^:* 0.35±0.06; *grh^IM^:* 0.37±0.06 clones) (Figure 1K). In contrast, a reduction was observed in the O-isoform specific *grh*^*370*^ with only 0.10±0.03 clones containing at least one ISC.

SInce *grh*^*370*^ clones at 10DPCI have fewer ISCs, we hypothesized that they would exhibit a decrease in proliferation relative to controls, which can be measured by counting the number of cells within a clone (22). Using clone size as a measure of ISC proliferation, we confirmed that the size of *grh*^*370*^ clones was significantly smaller, averaging 1 cell per clone compared to controls, with a mean of approximately 6 cells per clone (Figure 1L).

To distinguish the cell types that make up multicellular *grh* clones, the EC marker Pdm-1 and the EE marker Prospero (Pros) were assayed to determine the percentages of cells within clones with an EC (Pdm-1+ GFP+), EE (Pros+ GFP+) and ISC/EB fate (GFP+ only). The cellular composition between control and *grh* null clones (Figure 1M) was similar (ECs: 25-31%, EEs: 5-6% and ISC/EBs: 63-69%) suggesting that complete loss of Grh does not affect differentiation. The specific loss of all O-isoforms also did not prevent differentiation as *grh*^*370*^ cells positive for Pdm-1 (4% of *grh*^*370*^ cells) and Pros (9% of *grh*^*370*^ cells) could be detected, albeit Pdm-1 was detected at a much lower percentage than in control clones, however an increase in ISC/EBs (87% of *grh*^*370*^ cells) was observed. As loss of Grh O results in a loss of ISCs, it is likely that a majority of these GFP+ only cells are EBs. Taken together, these data imply that loss of O-isoform function results in differentiation of ISCs to EBs but also a reduced rate of EB to EC differentiation.

The increased severity of the *grh*^*370*^ MARCM clonal phenotype suggested that either loss of the O-isoform has a more severe effect than complete loss of Grh function, or that a second site mutation on the *grh*^*370*^ chromosome was responsible for the loss of ISCs. We therefore sought to validate the observed phenotype in two ways. Firstly, using Crispr methodology, we generated another *grh* mutation *(grh^WG^)* specific for O-isoform. Consistent with our analysis of *grh*^*370*^ clones, the number of *grh*^*WG*^ clones per midgut also decreased at 6DPCI (16.13 ± 4.55 clones) and 10DPCI (19.19 ± 3.44 clones) in comparison to controls (41.89 ± 3.54 and 44.96 ± 4.34, at 6DPCI and 10DPCI, respectively) (Figure 2A-C’). The number of cells per clone also significantly decreased in *grh*^*WG*^ clones compared to controls (Figure 2D-D’).

**Figure 2.**
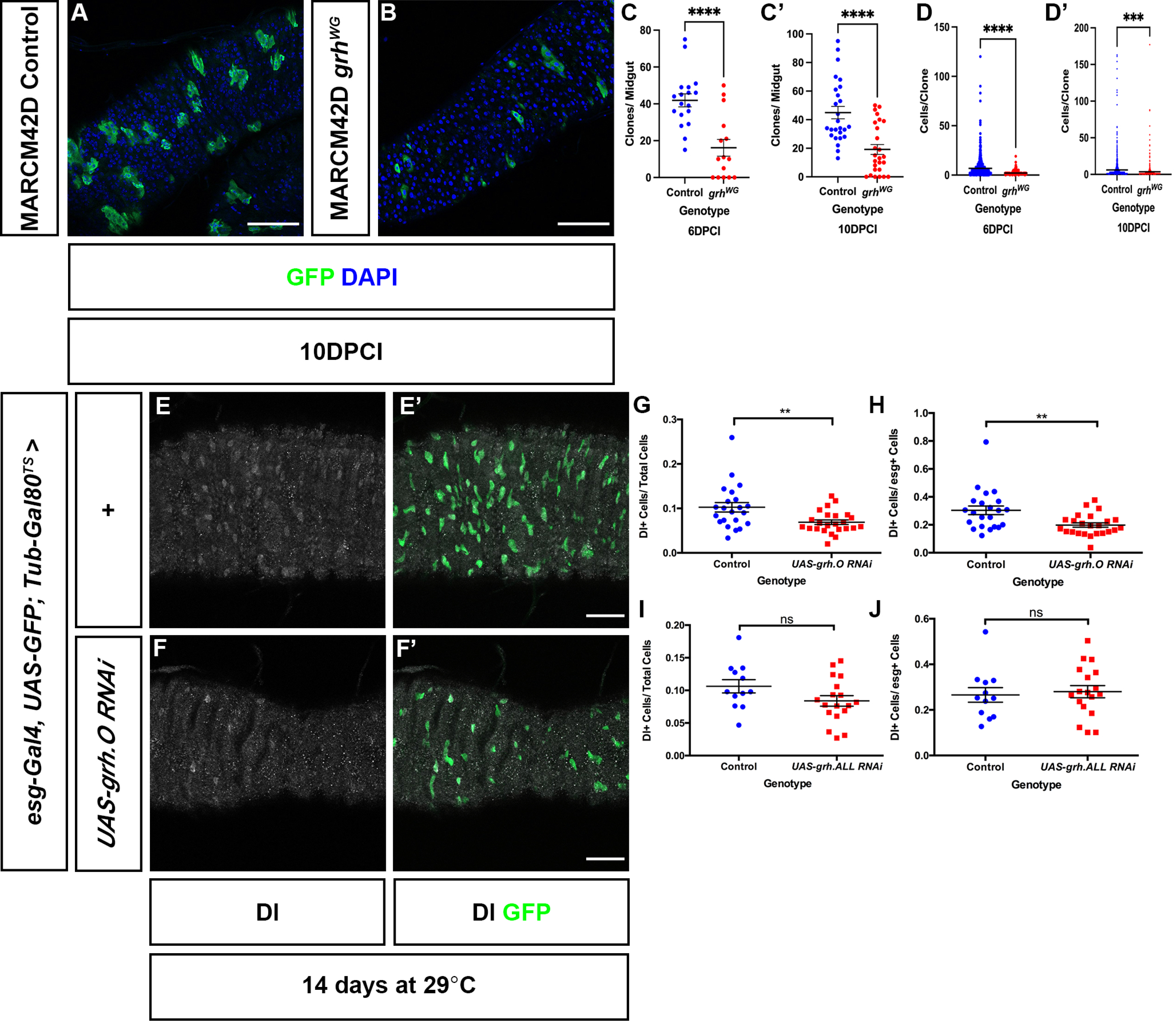
*grh*^*WG*^ MARCM Clones are also lost and O-isoform specific RNAi results in loss of ISCs. **A-B)** Confocal images of control **(A)** and *grh*^*WG*^ **(B)** MARCM clones maintained at 25C. Scale Bar 40μm. **C-C’)** Quantification of the number of clones per midgut in control (6DPCI: n= 18 midguts 10DPCI: n=26 midguts) and *grh*^*WG*^ (6DPCI: n=15 midguts, 10DPCI: n=26 midguts) MARCM clones at 6DPCI and 10DPCI. A reduction in the number of clones in *grh*^*WG*^ midguts is observed at both time points quantified. (Mean ± SEM, Unpaired Students T-test with Welsh’s correction, ****p<0.0001). **D-D’)** Quantification of clonal size between control and *grh*^*WG*^ MARCM clones. At both 6DPCI and 10DPCI, *grh*^*WG*^ clonal size, measured by counting the number of cells per clone is smaller than that of control clones. Control – 6DPCI: n=754 clones, 18 midguts; 10DPCI: n=1169 clones, 26 midguts. *grh*^*WG*^ – 6DPCI: n=242 clones, 15 midguts; 10DPCI: n=499 clones, 26 midguts. (Mean ± SEM, Unpaired Students T-test with Welsh’s correction, ****p<0.0001). **E-F)** Confocal images of control and *esg*^*TS*^> *UAS-grh.O RNAi* midguts. ISCs are marked by Dl and progenitor cells marked with GFP. Scale Bar 40μm. **G)** Quantification of Dl+ ISC cells in control (n=22 midguts) and *esg*^*TS*^> *UAS-grh.O RNAi* (n=24 midguts) midguts showed a decrease in the proportion of Dl+ cells over total cell number. (Mean ± SEM, Unpaired students T-test with Welsh’s Correction, **p=0.0078). **H)** Proportion of esg+ cells that are Dl+ ISCs in control (n=22 midguts) and *esg*^*TS*^> *UAS-grh.O RNAi* (n=24 midguts). (Mean ± SEM, Unpaired students T-test with Welsh’s Correction, **p=0.0049). **I)** Quantification of the proportion of Dl+ ISCs over total cell number in control (n=12 midguts) and *esg*^*TS*^>*UAS-grh.ALL RNAi* (n=18 midguts) show no significant difference between the two genotypes. (Mean ± SEM, Unpaired students T-test with Welsh’s Correction, ns = not significant). **J)** The proportion of esg+ cells that are Dl+ ISCs remain at similar levels in control (n=12 midguts) and *esg*^*TS*^> *UAS-grh.ALL RNAi* (n=18 midguts) midguts. (Mean ± SEM, Unpaired students T-test with Welsh’s Correction, ns = not significant).

As the phenotype recapitulated that observed with *grh*^*370*^, the loss of *grh*^*WG*^ in MARCM clones confirms that loss of Grh O-isoforms is responsible for the loss of ISCs and not a secondary unknown mutation in the *grh*^*370*^ background.

The requirement for Grh O-isoforms in maintaining ISCs was further confirmed via RNA interference. We generated a short hairpin RNA targeting exon 5 thus targeting O-isoforms. and expressed this *UAS-grh.O RNAi* transgene in ISCs and EBs using the *esg-Gal4, UAS-GFP; Tub-Gal80* (henceforth known as *esg*^*TS*^) driver. After 14 days at the permissive temperature, quantification of Dl+ cells in control midguts showed an average proportion of 0.10 Dl+ cells of the total cells. This measure significantly decreased to 0.07 in *esg*^*TS*^ > *UAS-grh.O RNAi* (Fig. 2G). Further analyses quantifying the ratio of ISCs to EBs (by examination of the proportion of Esg+ cells that were also Dl+) show a decrease of ISCs in *esg*^*TS*^ > *UAS grh.O RNAi* midguts containing a mean proportion of 0.20 Dl+ cells compared to an average of 0.30 Dl+ in controls (Figure 2H). This suggests that the majority of the relative proportion of GFP marked ISC/EBs (hereafter termed Esg+) in *esg*^*TS*^ > *UAS grh.O RNAi* midguts are EBs and that knock down of Grh O-isoforms results in premature differentiation of ISCs to EBs. No difference was observed using RNAi that targets all isoforms (Figure 2I-J), further indicating that it is the specific loss of Grh.O that results in loss of ISCs.

### Ectopic expression of N isoforms results in differentiation

The Grh O-isoform-specific phenotypes observed upon *grh*^*370*^ *and grh*^*WG*^ clonal analysis and in *esg*^*TS*^ > *UAS grh.O RNAi* midguts could be explained by an antagonistic interaction between the N- and O- isoforms in regulating ISCs. To test if increased N-isoform activity results in a reduced number of ISCs, Grh.N was over expressed in ISC/EBs using *esg^TS^.* After two days at the permissive temperature, there was a significant reduction in the relative proportion of Esg+ (or progenitor cells) in *esg*^*TS*^ > *UAS*-Grh.N midguts compared to controls (Figure 3A-B & E). Likewise, ectopic expression of another N-isoform, Grh.N’ (which contains a variant 3’ exon) also resulted in a loss of Esg+ cells (Figure 3F). Thus, ectopic expression of Grh N-isoforms in ISC/EBs results in the loss of ISC/EBs.

**Figure 3.**
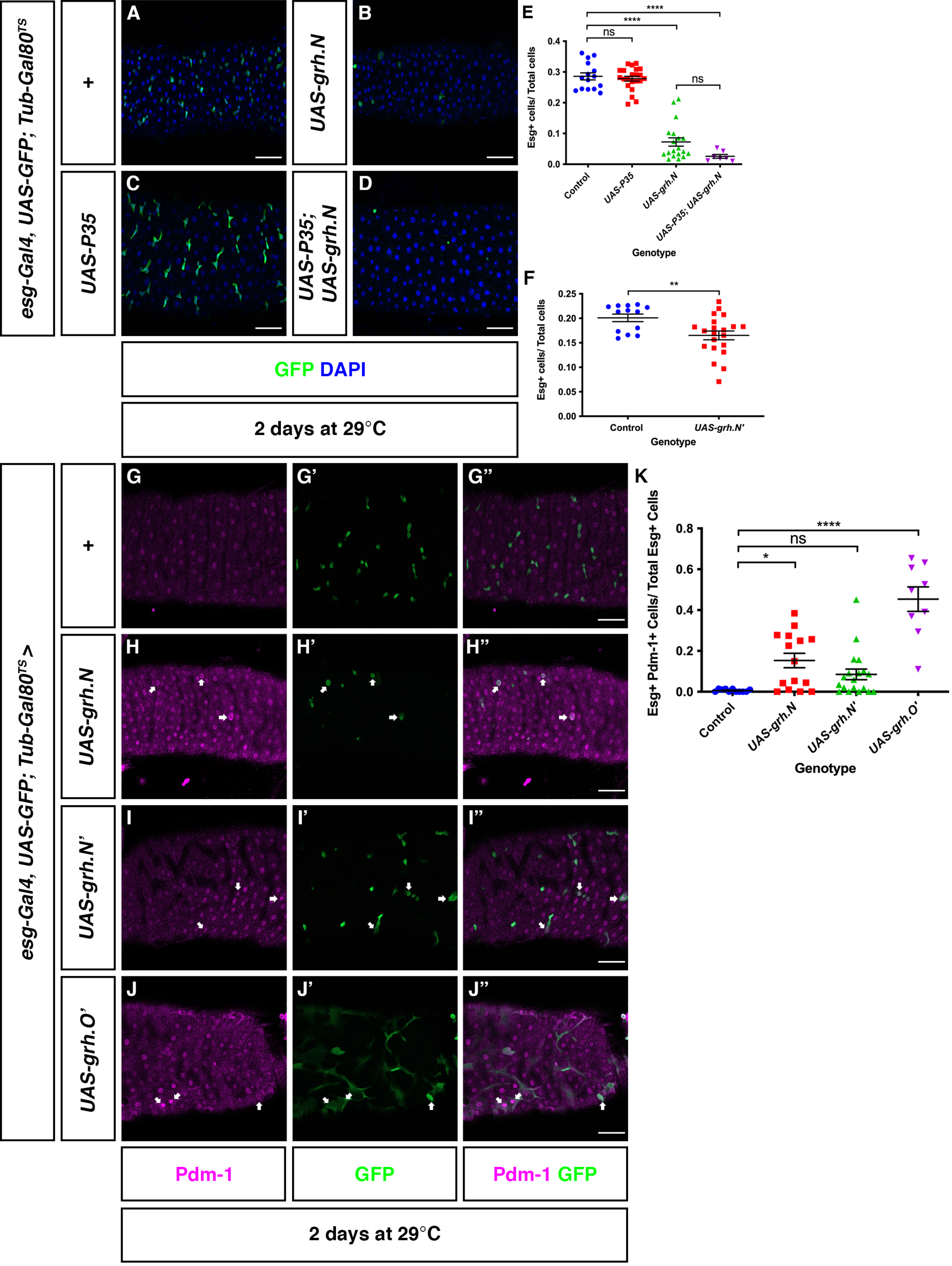
Ectopic expression of the N-isoform promotes differentiation. **A-D)** Representative images of control midguts **(A)**, midguts over expressing Grh.N **(B)**, the apoptotic inhibitor P35 **(C)** and midguts co-expressing P35 and Grh.N **(D)** in progenitor cells. Scale Bar 40μm. **E)** Quantification of the number of progenitor cells in control (n=15 midguts) midguts, midguts over expressing Grh.N (n=19 midguts), P35 (n=22 midguts) and midguts co-expressing Grh.N and P35 (n=7 midguts) shows that expression of p35 was not able to rescue loss of progenitor cells. (Mean ± SEM, One-Way ANOVA with Tukey’s Test, ns= not significant, ****p<0.0001). **F)** Quantification of the proportion of ISC/EBs in control and *esg*^*TS*^> *UAS-grh.N’* midguts. The proportion of progenitors in *esg*^*TS*^> *UAS-Grh.N’* midguts (n=21) have decreased in comparison to control midguts (n=13). (Mean ± SEM, Unpaired students T-test, **p=0.0088). **G-J)** Confocal images of control **(G)** and midguts ectopically expressing Grh.N **(H)** and Grh.N’ **(I)** and Grh.O’ **(J)** in ISC/EBs. Compared to controls, GFP+ progenitor cells expressing grh.N, Grh.N’ or Grh.O’ have increased cell size and also express the EC marker, Pdm-1 (arrows). Scale Bar 40μm. **K)** The proportion of esg+ progenitor cells expressing Pdm-1 in midguts increases in midguts over expressing Grh.N (n=15), Grh.N’ (n=9) and Grh.O’ (n=8) in comparison to control midguts (n=11). While the proportion of esg+ progenitor cells expressing Pdm-1 slightly increased in midguts overexpressing Grh.N’ (n=19), it was not statistically significant compared to controls. (Mean ± SEM, One-Way ANOVA with Tukey’s Test, ns=not significant, *p=0.0107, ****p<0.0001).

To investigate if the reduction of ISC/EBs due to the overexpression of Grh.N is a result of cell death, control and *esg*^*TS*^ > *UAS*- Grh.N midguts were immunostained with an antibody generated against human activated Caspase 3, which also detects apoptotic cells in *Drosophila* (23, 24). An activated Caspase 3 signal was not detected in control and *esg*^*TS*^ > *UAS*- Grh.N midguts suggesting that apoptosis is not the cause of ISC/EB loss (Supplemental Figure 1A-C). This was further confirmed by co-expressing the apoptotic inhibitor P35 (25) with *UAS*- Grh.N using the esg^TS^ driver (Figure 3C-E). Co-expression of P35 with Grh.N for two days at the permissive temperature was not sufficient in preventing the loss of Esg+ cells with the total proportion remaining at a similar level to the expression of Grh.N alone. The ISC/EB proportion in midguts only expressing P35 was comparable to controls demonstrating that expression of P35 does not affect ISC/EB number. Together these data illustrate that loss of ISC/EBs due to Grh.N overexpression is not a consequence of apoptosis.

The few Esg+ cells present that ectopically express Grh.N have large nuclei characteristic of ECs suggesting that they have differentiated (Figure 3H). In comparison to control midguts (Figure 3G), there was an increase in the proportion of Esg+ cells expressing Pdm-1 in midguts after ectopically expressing Grh.N in ISC/EBs (Figure 3K). While this trend was also observed when Grh.N’ was ectopically expressed using *esg*^*TS*^, quantification showed that it was not statistically significant. These data indicate that increased expression of at least one of the N-isoforms in ISC and EBs induces the process of differentiation down the EC lineage.

### *Grainy head* is expressed in the adult midgut

To determine if *grh* is expressed in the adult midgut, we designed primers to target subsets of *grh* mRNA transcripts (Supplemental Table 1). Additionally, primers for *esg*, an ISC/EB marker (9) and *sna*, a gene known to function in ISC and EBs (26) were used as positive controls. As expected, ddPCR (which allows absolute quantification of transcript levels) conducted on *w*^*1118*^ control midguts showed robust *esg* expression (Figure 4A) (1231 copies/μl ±318: Mean ± SEM) with expression of *sna* detectable at a much lower level (1.99 copies/μl ±0.96). Primers detecting all *grh* transcripts showed a an expression level of 5.60 copies/μl ±1.51, while transcripts responsible for the formation of O and O’-isoform expression were detected at a much lower level of 0.86 copies/μl ± 0.58.

**Figure 4.**
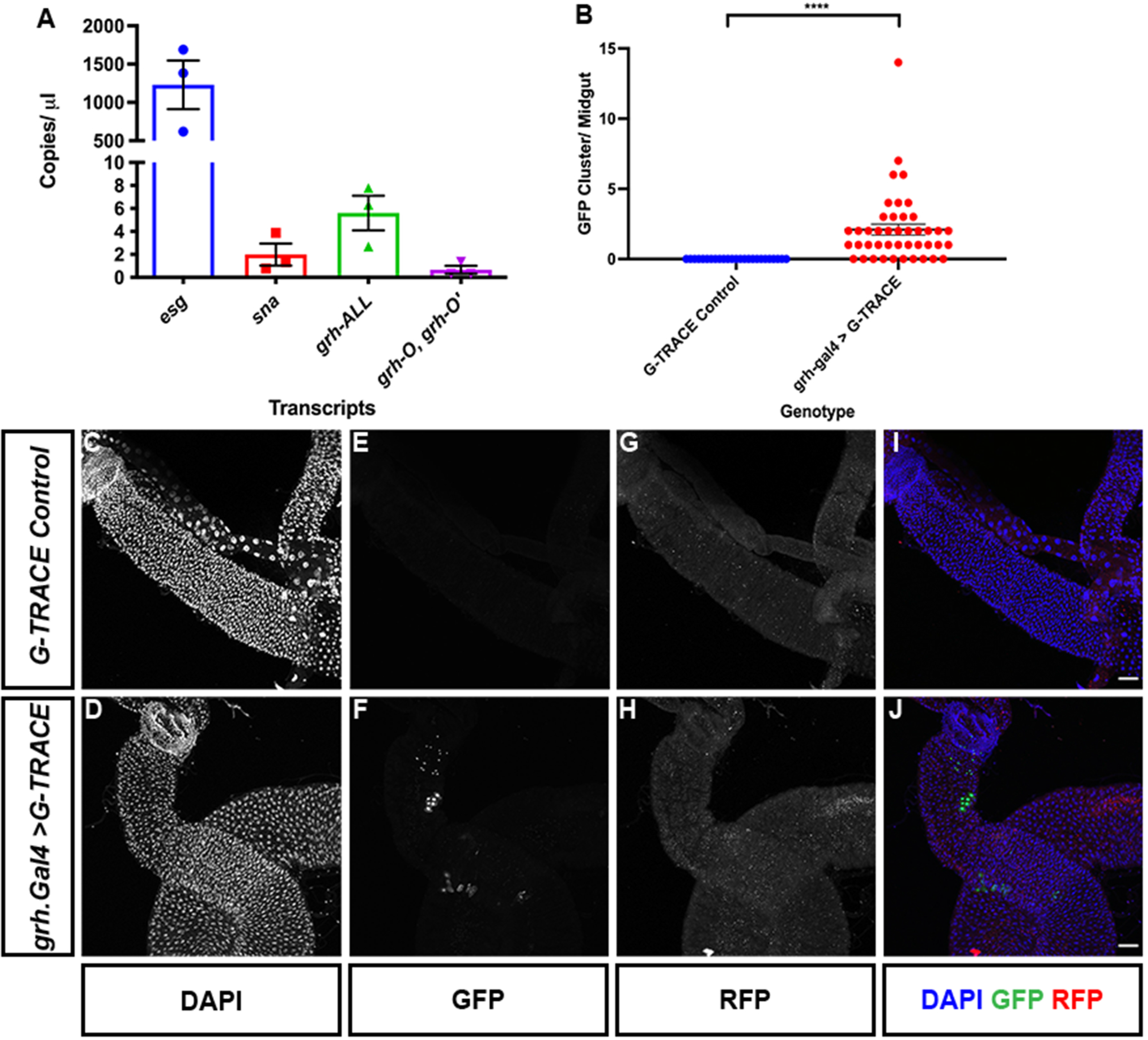
*grh* is expressed in the midgut. **A)** Quantification of *esg*, s*na* and *grh* mRNA transcripts in 5-7-day old adult *w*^*1118*^ midguts shows that in comparison to known ISC/EB marker *esg*, *grh* transcripts are expressed at much lower levels but is more comparable to that of ISC regulator, *sna.* mRNA quantification was conducted using biological triplicates (n=15 midguts for each replicate) with each replicate represented by individual data points. Bar graph represents mean ± SEM of data points. **B)** Quantification of the number of >2 cell clonal clusters per midgut in G-TRACE control and grh.Gal4>G-TRACE midguts. No clusters were observed in controls but grh.Gal4>G-TRACE midguts exhibited 2.09 ± 0.39 clusters per midgut. (Mean ± SEM, Unpaired students T-test with Welsh’s Correction, ****p<0.0001) **C-J)** Confocal images of G-TRACE only control **(C-I)** and midguts combining *grh^1249-G4^ (grh-Gal4)* with the G-TRACE reporter (w[*]; P{w[+mC]=UAS-RedStinger}4, P{w[+mC]=UAS-FLP.D}JD1, P{w[+mC]=Ubi-p63E(FRT.STOP)Stinger}9F6). No real time Gal4 activity was observed in either strain but compared to controls, GFP+ clonal clusters could be observed in grh.Gal4>G-TRACE. Scale Bar 40μm.

To investigate the spatial resolution of Grh, immunofluorescent staining of phenotypically wild-type midguts utilizing several Grh specific antibodies was performed. However, a signal could not be detected using four of the five antibodies (27–31) (Supplementary Figure 2A-D). While an immunofluorescent signal was detected using the final Grh antibody tested, this was later shown to be non-specific, with the signal remaining in GFP marked *grh* null mutant clones (Supplemental Figure 2E-F). While we were unable to detect Grh protein expression by antibody analysis, mutant analysis clearly indicates a role for Grh in the midgut. Therefore, we obtained a strain that contains a transposable element carrying the Gal4 open reading frame inserted into a Grh intron flanked by exons 9 and 10 of the Grh.RJ (Grh.O) transcript. This strain, *grh*^*1249-G4*^, expresses Gal4 under the influence of adjacent enhancer sequences (32). We crossed *grh*^*1249-G4*^ to a G-Trace (Gal4 Technique for Real time And Clonal Expression) line (33) that reports real time Gal4 activity via UAS-RFP and lineage traces cells that have expressed Gal4 via GFP. Grh enhancer activity in ISCs would therefore result in clusters of GFP labelled cells after shifting flies from 18C to 29C. We observed 2.09 ± 0.39 (Mean ± SEM) clusters of GFP+ cells per *grh*^*1249-G4*^>G-Trace midgut but none in G-Trace alone midguts after 5 days at 29C (Figure 4B-4J). These data suggest that Grh transcripts are present in the midgut and that Grh transcription is active in ISCs and possibly EBs.

### Ectopic Grh O-isoforms promote a delay in differentiation that results in cells having characteristics of EBs and ECs

To examine the effect of elevating O-isoform levels, we ectopically expressed Grh.O’ in ISC/EBs using the *esg*^*TS*^ driver. Increased expression of Grh.O’ resulted in an increase in the proportion of Esg+ cells when compared to controls (Figure 5A-C). This could represent an increase in either ISCs or EBs. To differentiate the two cell types, control and *esg*^*TS*^ > *UAS*- Grh.O’ midguts were immunostained with Dl, with analysis revealing that the majority of Esg+ cells were Dl positive, suggesting an increase in ISCs (Figure 5A-B). Interestingly, cells labelled with both Dl and GFP in *esg*^*TS*^ > *UAS*- Grh.O’ midguts appeared enlarged (compared to ISCs), a property of ECs which undergo endoreplication and an increase in cellular size. Labelling control and *esg*^*TS*^ > *UAS*- Grh.O’ midguts with Pdm-1 revealed that co-expression of GFP (Esg+) with Pdm-1 was commonly observed in *esg*^*TS*^ > *UAS*-Grh.O’ midguts, but not control midguts (Figure 3J-K). The expression of Pdm-1 in GFP positive cells in *esg^TS^ >UAS*-Grh.O’ cells indicates that increased Grh.O’ activity also induces differentiation, although the maintenance of Dl expression shows the cells retain some ISC properties.

**Figure 5.**
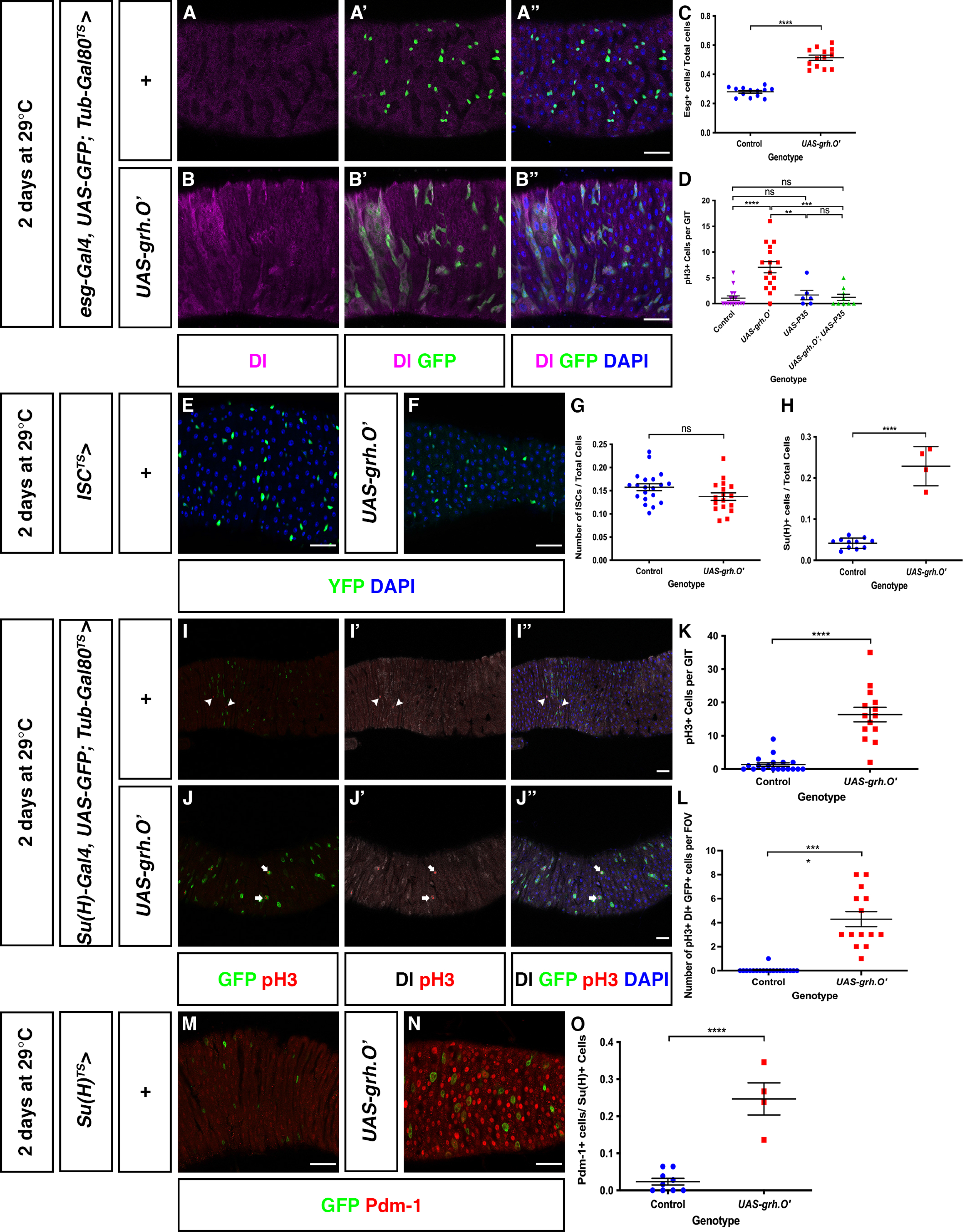
Overexpression of O-isoform results in both ISC maintenance and differentiation. **A-B)** Control midguts and midguts over expressing Grh.O’ in progenitor cells. While only some progenitor cells express ISC marker, Dl **(A)** a majority of *esg*^*TS*^ > UAS-Grh.O’ cells express Dl **(B)**. **C)** In comparison to control midguts (n=13) there is an increase in the proportion of esg+ cells in midguts over expressing Grh.O’ (n=13). (Mean ± SEM, Unpaired students T-test with Welsh’s Correction, ****p<0.0001). **D)** An increase in the mitotic index as measured by the number of pH3+ cells are observed in midguts ectopically expressing Grh.O’ (n= 16). This increase in mitotic index is rescued by the co-expression of the apoptotic inhibitor p35 with Grh.O’ (n= 9) with its mitotic index returning to levels seen in control midguts (n= 16) and midguts only expressing p35 (n= 6). (Mean ± SEM, One-Way ANOVA with Tukey’s Test, **p=0.0027, ***p=0.0001, ****p<0.0001). **E-F)** Confocal images of control midguts and midguts ectopically expressing Grh.O’ in only ISCs (green). Ectopically expressing Grh.O’ in ISCs does not appear induce differentiation with the YFP+ ISCs remaining at a similar size to YFP+ cells in control midguts. Scale Bar 40μm. **G)** The proportion of ISCs in control midguts (n=20) and midguts ectopically expressing Grh.O’ (n=17) remain at similar level. (Mean ± SEM, Unpaired students T-test with Welsh’s Correction, p=0.0749). **H)** In comparison to controls (n=11 midguts), an increase in GFP+ EBs (express Su(H) EB marker) is observed in midguts ectopically expressing Grh.O’ (n=4). (Mean ± SEM, Unpaired students T-test with Welsh’s Correction, ****p<0.0001). **I-J)** Representative images of control midguts **(I)** and midguts over expressing Grh.O’ **(J)** in EBs (driven from Su(H)Gal4). In control midguts, ISCs are solely marked by Dl with mitotically active ISCs marked by pH3 (arrowheads) whereas overexpression of Grh.O’ in EBs induces EB expression of Dl and EB mitotic division (arrows). Scale Bar 40μm. **K)** Comparison of the mitotic index in control midguts (n=19) and midguts over expressing Grh.O’ (n=14) in EBs. An increase in the mitotic index is observed in midguts over expressing Grh.O’ in EBs. (Mean ± SEM, Unpaired students T-test, ****p<0.0001). **L)** Quantification of the number of EBs undergoing mitotic division. (Mean ± SEM, Unpaired students T-test, ****p<0.0001). **M-N)** Confocal images of control midguts and midguts over expressing Grh.O’ in EBs immunostained with EC marker, Pdm-1. Whereas GFP+ EBs in controls rarely express Pdm-1, an increase in *Su(H)*^*TS*^ > UAS-Grh.O’ cells are positive for Pdm-1 is observed. Scale Bar 40μm. **O)** The proportion of EBs expressing Pdm-1 increases in *Su(H)*^*TS*^ > UAS-Grh.O’ midguts (n=4) in comparison to control midguts (n= 11). (Mean ± SEM, Unpaired students T-test, ****p<0.0001).

The accumulation of progenitor cells with confused identity is reminiscent to the age-related decline in epithelial structure that leads to cell death (34). Typically, the induction of apoptosis in the midgut leads to compensatory ISC proliferation (35). This was indeed observed in midguts over-expressing Grh.O’ with the mitotic index, measured by counting the number of Phospho-histone 3 positive cells, increasing to a mean of 7 compared to a control mean of 1 (Figure 5D). Moreover, the increase in mitotic index was rescued (returning to a mean of ~1) when apoptotic inhibitor P35 was co-expressed with Grh.O’. These data suggest that overexpression of Grh.O’ in ISC and EBs results in cells with confused identity and ultimately cell death. Consequently, it is EB/EC cell death that stimulates ISC proliferation.

Our data suggest that overexpression of Grh.O’ can induce both differentiation and maintenance of ISC identity. The results presented in Figure 1M suggest that the O-isoform is required to prevent ISC differentiation to EBs, but also to enhance EB to EC differentiation. In order to determine the effect of Grh.O’ overexpression in ISCs alone, we utilised *esg-Gal4, UAS-YFP; Su(H)-Gal80, Tub-Gal80*^*TS*^ (hereafter termed *ISC*^*TS*^) which will inhibit Gal4 mediated expression in EBs. Ectopic expression of Grh.O’ in ISCs did not result in an increase in ISC number with the number of ISCs remaining at an equivalent level to that of controls (Figure 5G). Moreover, the increase in Grh.O’ expression did not cause premature differentiation into ECs as GFP+ nuclear size remained at a similar size to that of controls (Figure 5E-F).

The function of Grh.O’ in EBs was similarly investigated by using *Su(H)GBE-Gal4, UAS-GFP; Tub-Gal80*^*TS*^ (hereafter termed *Su(H)*^*TS*^). Ectopic expression of Grh.O’ in EBs led to an increase in the proportion of GFP+ EBs in comparison to controls (Figure 5H). The increase in EB number appears to be caused by an increase in the mitotic index of *Su(H)*^*TS*^ > *UAS*-Grh.O’ midguts (Figure 5I-K). Somewhat surprisingly, a number of these mitotically active cells were GFP+ suggesting that ectopic expression of Grh.O’ in EBs may facilitate EB cell division (Figure 5J & L). Given that cell division is characteristic of ISCs, it is possible that these cells may have some ISC-like identity.

In addition to maintaining ISC identity in *esg*^*TS*^> Grh.O’ midguts, overexpression of Grh.O’ was also able to induce differentiation into the EC lineage. This ability was also investigated in EBs. Analysis of *Su(H)*^*TS*^ > Grh.O’ midguts showed that there was an increase in the number of EBs expressing the EC marker, Pdm-1 (Figure 5M-O) suggesting that Grh.O’ is also able to induce EB differentiation to ECs.

## Discussion

Here we have provided evidence that Grh is required to maintain ISCs in the *Drosophila* midgut. Surprisingly, it is the function of the O-isoforms that is crucial for ISC maintenance. Generation of *grh*^*370*^ midgut clones, including a new CRISPR-generated allele, demonstrated that ISCs that do not express an O-isoform are lost from the epithelium.

It is unusual for a hypomorphic mutation to exhibit a more severe phenotype than a null but this is not unprecedented and can result from different products of a gene acting in opposing manners. For example, the murine *Trp73* gene encodes two major classes of isoforms, resulting from alternative initiation sites, termed TAp73 and ΔNp73 (36). Each isoform has unique properties and it appears to be the balance of isoforms in a cell that determines phenotypic outcome. Specific isoform knockouts in mice exhibits unique phenotypes. *TAp73*^−/−^ mice develop tumours not observed in *Trp73*^−/−^ nulls (37).

If an antagonistic relationship exists between Grh O- and N-isoforms it would be expected that overexpression of an N-isoform may generate a similar phenotype to loss of the O-isoform. Overexpression of an N-isoform resulted in a loss of ISC and EBs via forced differentiation. However, differentiation proceeded in *grh* null clones. This suggests that N-isoforms, while they may facilitate differentiation, are not critical in the absence of O-isoforms and other factors may compensate for their loss.

Overexpression of the O’-isoform in ISCs/EBs resulted in cells with confused identities. While a majority of the Esg+ cells expressed the ISC marker, Dl, they also exhibited increased cell size and expressed the EC marker, Pdm-1. This suggested a role for O-isoforms in both ISC maintenance and EC differentiation. It is possible that the high level of ectopic expression produced by GAL4 overrides the nature of the isoform being expressed but the phenotype observed with ectopic Grh.O’ was qualitatively different to that from Grh.N (or Grh.N’).

The dual role of Grh.O was also observed when this isoform was specifically expressed in EBs as it resulted in cells that express both Pdm-1 and Dl. Interestingly, overexpression of Grh.O in only ISCs did not induce an increase in ISC number suggesting that it alone cannot specify ISC identity. *grh*^*370*^ clones lose stem cells but increase the proportion of GFP+ Pdm-1- Pros-cells compared with control clones suggesting that they have an increased proportion of EBs, or that ISCs differentiate rather than being lost via apoptosis. These data imply that Grh O-isoforms maintain ISCs by preventing them from prematurely differentiating. The data also show a decrease in the proportion of GFP+ Pdm-1+ cells in *grh*^*370*^ clones, suggesting that the accumulated EBs have an impaired ability to differentiate into ECs, again suggesting a dual role for Grh O-isoforms to 1) maintain the ISC state, and 2) to facilitate EB to EC differentiation.

A reiterative, but different, function for Grh O-isoforms in two stages of the ISC to EC differentiation process may also explain why *grh* is expressed at such low levels in the midgut. A very tight control of *grh* expression would permit function in ISCs, which may be accompanied by a transient reduction to permit EB differentiation and subsequent upregulation to allow differentiation to ECs. It is likely that a network of transcriptional regulators act to facilitate midgut differentiation that may be at, or beyond, the resolution of detection by current RNAseq pipelines. This underscores the importance of genetic screens that may be capable of detecting phenotypes associated with genes that could be potentially ignored from transcriptome analyses.

The mechanism of how Grh.O exerts its specificity is yet to be determined. The O-specific sequences do not appear to be conserved in vertebrate GRHL proteins despite them exhibiting differential splicing (38), however differential activities of the vertebrate proteins offer the suggestion that isoform specific functions observed in *Drosophila* have evolved into separate gene functions in vertebrates. This is exemplified in studies that have examined GRHL function in tumorigenesis. Depending on the cancer type, GRHL proteins appear to have oncogenic or tumour suppressive roles. Quan, Jin (39) and Quan, Xu (40) showed GRHL2 upregulation in colorectal cancer samples that correlated with tumor progression and advanced clinical stage. This oncogenic role is underlined by functional studies overexpressing GRHL2 in CRC cells lines, which induced cellular proliferation (40). Converse to the oncogenic function of GRHL2, GRHL3 can act as a tumour suppressor as loss of GRHL3 in keratinocytes suppresses Pten expression leading to subsequent activation of PI3K/AKT/mTOR signaling resulting in squamous cell carcinoma (41). These examples of differential effects of GRHL proteins upon tumour formation and progression may be associated with different contexts of GRHL expression or may represent divergence of GRHL function, similarly to the *Drosophila* Grh isoforms. In either case it is likely that the GRHL proteins are interacting with different cofactors to exert different effects upon tumourigenesis. Further work will determine how GRH/GRHL proteins and isoforms co-operate in regulating epithelial stem cell maintenance and differentiation.

## Materials and Methods

### *Drosophila* Stocks and Husbandry

Fly stocks were maintained on a standard culture medium at 25°C unless otherwise specified. Mated female flies were exclusively analysed throughout this study, due to differences in male and female ISC behavior (42). A detailed list of fly strains used in this study are listed in Supplementary Table 2. Flies containing the *Tub-Gal80*^*TS*^ allele were crossed and maintained at the non-permissive temperature of 18°C. Following eclosion, 3-5-day old flies were transferred to the permissive temperature of 29°C for further analysis.

### Generation of Marked Clones

The Mosaic Analysis with Repressible Cell Marker System (MARCM) was used to generate positively marked GFP homozygous clones. Unless otherwise stated, MARCM crosses were established and maintained at 18°C. 3-5 day old adult flies of the genotypes *UAS-cd8 GFP, hs-FLP/ +; frt42DtubGal80/ frt42D; tub-Gal4/+, UAS-cd8 GFP, hs-FLP/ +; frt42DtubGal80/frt42Dgrh^S2140^; tub-Gal4/+, UAS-cd8 GFP, hs-FLP/ +; frt42DtubGal80/ frt42Dgrh^IM^; tub-Gal4/+, UAS-cd8 GFP, hs-FLP/ +; frt42DtubGal80/ frt42Dgrh^370^; tub-Gal4/+* were heat shocked for 1 hour in a 37°C running water bath. Flies were then returned to 18°C to minimize the incidence of leaky MARCM clones. Intestines were analyzed 5-10 days after clonal induction.

### Generation of *UAS-grh.O* RNAi Line

Short hairpin design was conducted using *grh* exon 5 sequence with top candidate selected using DSIR website (43). Selected hairpin, 5’-CGGGATCAGACAAATATCCAA-3’ was cloned into pWALIUM20 (44) with potential plasmids verified by Sanger sequencing. Verified plasmid was then injected into embryos (BestGene Inc. Co) and a stable transgenic line made.

### Generation of *grh*^*WG*^ allele

CRISPR mediated mutagenesis, performed by WellGenetics Inc. was used to insert 3-frame stop codons in *grh* exon 5 to generate a C-terminal truncation only affecting O-isoforms (*grh-RJ, RL, RN and RO)*. Briefly, DNA plasmids containing hs-Cas9, *grh* gRNAs targeting exon 5 and a cassette containing 2 loxP sites, selection marker 3xP3-RFP, two homology arms and 3-frame stop codons were injected into *w*^*1118*^ embryos. F1 flies carrying 3xP3-RFP selection marker were then validated by PCR, sequencing and backcrossing to *grh*^*IM*^ and *Df(grh)*.

### G-Trace analysis

The Grh enhancer trap, *grh*^*1249-G4*^, was crossed to the G-Trace (BL2820), w[*]; P{w[+mC]=UAS-RedStinger}4, P{w[+mC]=UAS-FLP.D}JD1, P{w[+mC]=Ubi-p63E(FRT.STOP)Stinger}9F6/CyO, strain to allow lineage tracing of cells that have expressed Gal4 via UAS-FLP recombinase combined with Ubiquitin-p63 promoter upstream of a STOP sequence flanker by FRT sites and GFP. Gal4 activity will result in removal of the stop sequence and expression of GFP in the cell that saw Gal4 activity plus all subsequent daughter cells. Real time Gal4 activity was also observed via UAS-RFP (although no expression was detected under our conditions of raising flies at 18C followed by a shift to 29C for 5 days to permit optimal Gal4 activation).

### Immunostaining

Adult mated female *Drosophila* midguts were dissected in Phosphate Buffered Saline (PBSx1). Dissected midguts were then fixed in 4% formaldehyde for 1 hour or overnight at 4°C. Following fixation, midguts were then washed 3x in PBT (PBS + 0.1% Triton-X100) for 5 minutes each and blocked for 1 hour in PBTH (PBS + 0.1% Triton X100 + 5% Horse Serum). Midguts were then incubated in the following primary antibodies: chicken anti-GFP (1:2000, AbCam), mouse anti-Delta (1:100, DSHB), rabbit anti-Pdm-1 (1:2000, Xiaohang Yang), rabbit anti-pH3 (1:5000, Upstate), mouse anti-β-Galactosidase (1: 20, DSHB), chicken anti-β-Galactosidase (1:2000, AbCam), rabbit anti-β-Galactosidase (1:2000, Cappel), mouse anti-GRH (1:5, Sarah Bray), rat anti-GRH (1:500, Stefan Thor), rabbit anti-GRH (1:200, William McGinnis /1:1000, Melissa Harrison), rabbit anti-Activated Caspase 3 (D175) (1: 200, Cell Signaling Technologies) overnight at 4°C. After primary antibody incubation, the samples were washed 3x for 5 minutes in PBT and incubated in corresponding Alexa Fluor (1:500, Invitrogen) secondary antibodies for 2 hours. This was then followed by a DAPI wash for 20 minutes and 3x washes in PBT for 5 minutes each. Samples were then mounted in 80% glycerol. All steps were carried out at room temperature unless otherwise stated.

### Image Acquisition and Analysis

Images were acquired on the Zeiss LSM800 or LSM880 Confocal Microscopes as serial optical sections (z-stacks) of 1,024 x 1,024 resolution. Only the first 500μm adjacent to the pyloric ring was imaged and analysed throughout this study. This was to ensure consistent comparison of regions between midgut samples and the prevention of confounding results due to regional midgut differences (45, 46).

Clonal cell counts were conducted using Bitplane Imaris software to render 3D reconstructions of intestines. Imaris Spot and Surface functions were then utilized to quantify and map cell nuclei to clones.

Cell counts and were performed using FIJI/ ImageJ software. FIJI was also used to process images and Adobe Photoshop used to compile figure panels. Statistical analysis and graphs were created using Graphpad PRISM.

### Droplet Digital PCR

Droplet Digital PCR (ddPCR) was used to for expression analysis during this study. Taqman gene expression assays were purchased from Thermofischer Scientific (Supplementary Table 1). RNA was extracted from at least 15 midguts per genotype using the Qiagen RNeasy Mini Kit (Cat no. 74104). RNA quality was analysed on the Agilent Tapestation 2200. RNA with an RNA Integrity Number (RIN) above 8 was used for analysis. RNA quantification was carried out on a Qubit 4 Fluorometer. 500ng of RNA was then used for cDNA synthesis using the Bioline Sensifast cDNA Synthesis Kit (Cat no. BIO-65053) as per the manufacturer’s instructions.

## Supporting information

Supplemental data, methods and references

## Acknowledgements

The authors would like to thank the Bloomington *Drosophila* Stock Center, Vienna *Drosophila* RNAi Center, Kyoto *Drosophila* Stock Center, Australian *Drosophila* Biomedical Research Support Facility (OzDros), University of Melbourne Biological Optical Microscopy Platform, S.Hou, S. Bray, K. Harvey, L. Quinn, A. Gould, C. Samakovlis and DL. Jones for *Drosophila* strains; the Developmental Studies Hybridoma Bank, W. McGinnis, M.Harrison, S. Bray, and S. Thor for antibodies.

## Author Contributions

ND, JH and FC conducted all experiments. ND, NAS and GRH planned experiments. All authors were involved in writing and editing the manuscript.

## Ethics Statement

Use of genetically modified Drosophila was approved by the University of Melbourne Gene Technology and Biosafety Committee (IBC reference no. 2022/014 and 2017/023)

## Data availability

All data used for this study and not presented in the text or figures are available upon request. Transgenic *Drosophila* lines generated for this study are available upon request.

