## Supplemental data, methods and references for "Alternate Grainy head isoforms regulate *Drosophila* midgut intestinal stem cell differentiation"

**Supplemental figures, figure legends and tables for Dominado et. al.**

Supplemental Figure 1

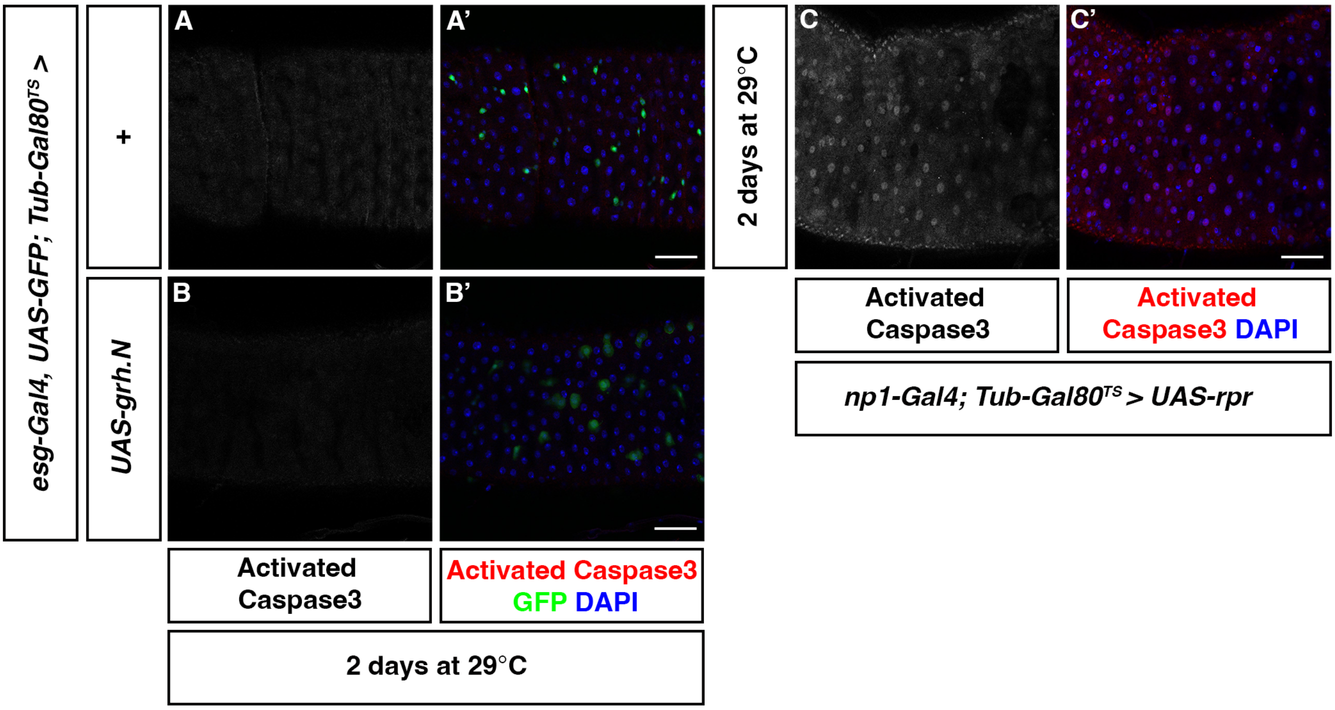

Supplemental Figure 2

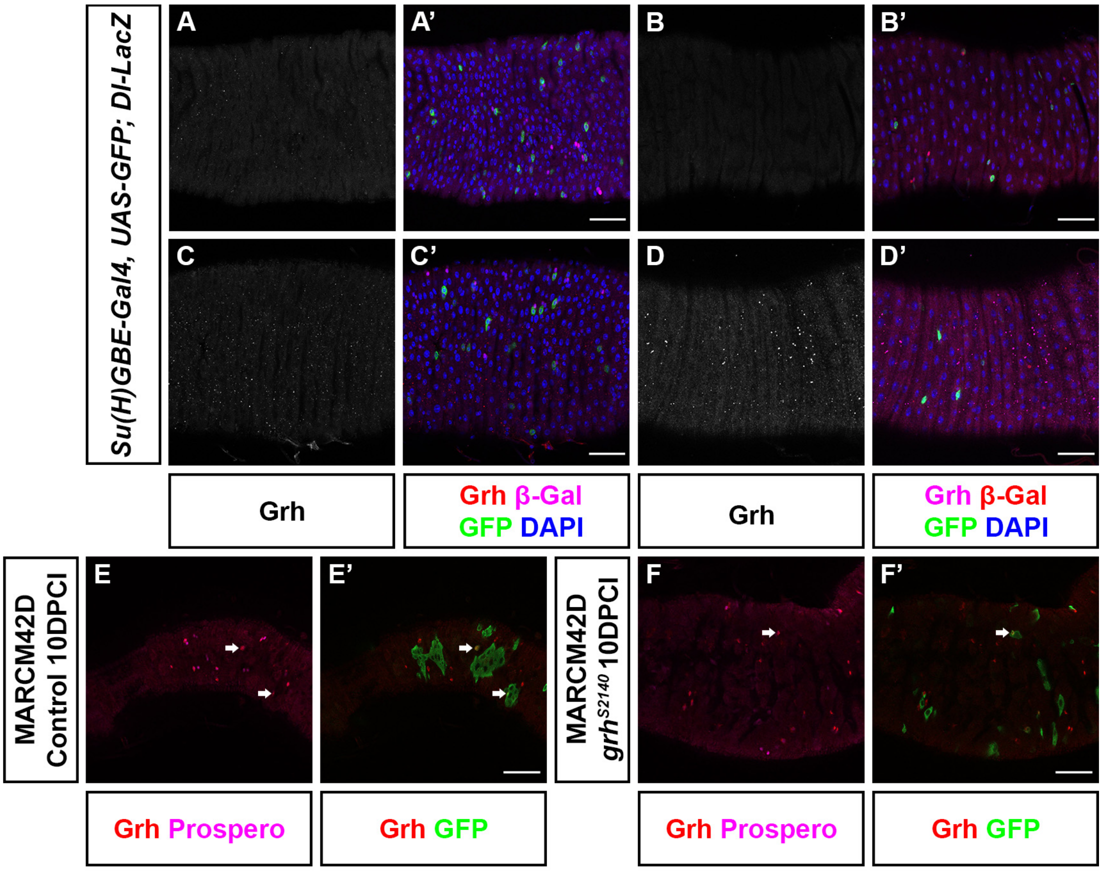

**Supplemental Figure Legends**

**Supplemental Figure 1. Overexpression of Grh.N does not cause cell death**

**A-B)** Confocal images of control midguts **(A)** and midguts over expressing N-isoform **(B)** in ISC/EBs. Activated caspase 3 was not detected in control and *esg*$^{TS}$> *UAS*-Grh.N midguts. Scale Bar 40μm.

**C)** Representative image of midgut expressing cell death gene, *reaper* in ECs and immunostained with activated caspase 3 for a positive control. Scale Bar 40μm.

**Supplemental Figure 2. Grh protein expression could not be discerned using antibodies**

**(A-D)** Adult midguts immunostained with four different Grh antibodies. Grh staining was not observed in phenotypically wild type midguts. ISCs were labelled by the reporter *Dl-LacZ* while *Su(H)GBE-Gal4>UAS-GFP* was used to mark EBs. Scale Bar 40μm.

**(E-F)** A fifth Grh antibody detects a cytoplasmic Grh (red) signal in the midgut that co-localizes with Prospero positive EE cells (magenta). However, this signal is not specific to Grh in the midgut as it can be detected in both control and homozygous *grh* null mutant MARCM clones (green). Scale Bar 40μm.

**Supplemental Table 1: TaqMan Gene Expression Assays**

| Gene Name | Assay ID |
|---|---|
| *sna* | Dm01841564_s1 |
| *esg* | Dm01841264_s1 |
| *grh-ALL* | Dm01816396_m1 |
| *grh-O, O'* | Dm01841950_m1 |
| *RpL32* | Dm02151827_g1 |
| *RpL11* | Dm01842483_g1 |

**Supplemental Table 2: Fly Strains Used in this Study**

| Fly Strain | Source |
|---|---|
| $w^{1118}$ | Bloomington Drosophila Stock Centre BDSC_3605 |
| $grh^{S2140}$*/CyO* | BDSC_10460 |
| *Su(H)GBE-Gal4, UAS-GFP / CyO* | Stephen Hou |
| *Delta-LacZ* | BDSC_11651 |
| $grh^{06850}$*/CyO* | BDSC_12325 |
| $esg^{TS}$ *(esgGal4, UAS GFP; TubGal80*$^{TS}$*/ [SM6B-TM6B])* | Kieran Harvey |
| *MARCM42D (hs-FLP, UAS-CD8GFP; FRT42DtubGal80/ CyO; Tub-Gal4/TM6B)* | Leonie Quinn |
| *b pr cn grh*$^{370}$*bw/CyO* | Alex Gould |
| $cn^1$ $grh^{IM}$ $bw^1$*/ SM6a* | BDSC_3270 |
| *UAS-grh.O'* | BDSC_42227/42228 |
| *UAS-grh.N* | Christos Samakovlis |
| *esg;Su(H)::Gal80 (w; esg-Gal4, UAS-2xYFP/ CyO; Su(H)Gal80, TubGal80*$^{TS}$*/ TM3)* | Leanne Jones |
| *UAS-grh.N'* | This Study |
| *UAS-10x-UAS-IVS-myr-GFP* | BDSC_32197 |
| *UAS-grh.O RNAi* | This Study |
| *UAS-P35* | BDSC_5073 |
| $grh^{WG}$*/ CyO* | This Study |
| $grh^{1249\text{-}G4}$ *(grh-Gal4)* | BDSC_65637 |
